# Computer vision for assessing species color pattern variation from web-based community science images

**DOI:** 10.1101/2022.02.11.480114

**Authors:** Maggie M. Hantak, Robert P. Guralnick, Alina Zare, Brian J. Stucky

**Affiliations:** Florida Museum of Natural History, University of Florida, Gainesville, FL 32611, USA; Department of Electrical, and Computer Engineering, University of Florida, Gainesville, FL 32611 USA; Agricultural Research Service, U.S. Department of Agriculture, Beltsville, MD 20705, USA

## Abstract

Openly available community science digital vouchers provide a wealth of data to study phenotypic change across space and time. However, extracting phenotypic data from these resources requires significant human effort. Here, we demonstrate a workflow and computer vision model for automatically categorizing species color pattern from community science images. Our work is focused on documenting the striped/unstriped color polymorphism in the Eastern Red-backed Salamander (*Plethodon cinereus*). We used an ensemble convolutional neural network model to analyze this polymorphism in 20,318 iNaturalist images. Our model was highly accurate (∼98%) despite image heterogeneity. We used the resulting annotations to document extensive niche overlap between morphs, but wider niche breadth for striped morphs at the range-wide scale. Our work showcases key design principles for using machine learning with heterogeneous community science image data to address questions at an unprecedented scale.

## Introduction

Species color patterns represent model systems for understanding evolution because color is a quantifiable biological trait that provides pertinent information about the organism. For instance, color patterns are used as a signal in mate choice and predator-prey interactions, and can aid in thermoregulation (Endler and Mappes, 2017). Color polymorphic species, in which multiple phenotypes (i.e., color morphs) coexist within the same population (Ford, 1945), make particularly good models for studying evolutionary change, as color patterns are discrete, and color morph frequency often varies geographically (McLean and Stuart-Fox, 2014). Further, morphs comprise correlated trait complexes, resulting in divergent selective pressures for a single species (Sinervo and Svenson, 2002; Mckinnon and Pierotti, 2010).

A wealth of information regarding species color patterns exists in web-based community science platforms, in which contributors can upload their own photographs of animals and plants, and seek help from other participants in identifying their observations. One of the largest and most successful platforms is iNaturalist (http://www.inaturalist.org/), which as of January 2022, holds > 88 million images of various species from across the world and roughly doubles in size each year. DiCecco et al. (2021) showcase the research value of iNaturalist, but one still nascent application is broad-scale assembly of color pattern data (but see Lehtinen et al., 2020; Lattanzio and Buontempo, 2021). The key challenge is that manual extraction of color pattern data is time and effort intensive. Automation is an obvious next step but complex image backgrounds can confuse simplistic image analysis toolkits (Peña et al., 2014; Pollicelli et al., 2020). Therefore, developing best practices and tools for streamlining extraction of information from variable quality images submitted by amateur naturalists is a critical need for processing the plethora of digital image data now being generated, enabling data-intensive research efforts in the areas of ecology and evolutionary biology (Weinstein, 2018; Lürig et al., 2021).

Artificial intelligence methods, and deep learning in particular, offer the most promise for automating collection of phenotypic data (Lürig et al., 2021), given their remarkable ability to make accurate predictions. Convolutional neural networks (CNNs) are the basis for current state-of-the-art accuracy in whole image classification (Deng et al., 2009; Zeiler, 2014; Sermanet et al., 2014). A CNN is a deep learning algorithm that uses training data to learn how to extract features from input images and then use those features to interpret an image’s content (LeCun et al., 2015). Much recent work using CNNs for ecological studies has focused on species identification from complex images (e.g., camera-trap images; Wäldchen and Mäder, 2018; Tabak et al., 2019; Willi et al., 2019; Whytock et al., 2021). Less developed are deep learning approaches that score quantitative traits of interest on those images.

Here, we present a workflow and machine learning approach for classifying color patterns of animals from community science photographs. To illustrate the value of this computer vision model, we focus on a use-case of a striped/unstriped color pattern polymorphism in the geographically widespread and abundant Eastern Red-backed Salamander, *Plethodon cinereus* (Petranka, 1998). The ‘striped’ color morph exhibits a stripe that varies in color from yellow to dark red, which is overlaid on a black dorsum, and the ‘unstriped’ morph is completely dark in dorsal coloration (Fig. 1). The ecological and evolutionary mechanisms influencing the geographic patterns of coloration in *P. cinereus* color morphs remains unclear, and little work has been done to examine range-wide patterns of the polymorphism (but see Gibbs and Karraker, 2006; Moore and Ouellet, 2015; Cosentino et al., 2017). Studies from single populations have suggested that the color morphs are correlated with distinct climatic niches; the striped morph is more associated with cooler, wetter niches, while the unstriped morph is more associated with warmer, drier conditions (Moreno, 1989; Anthony et al., 2008).

**Figure 1.**
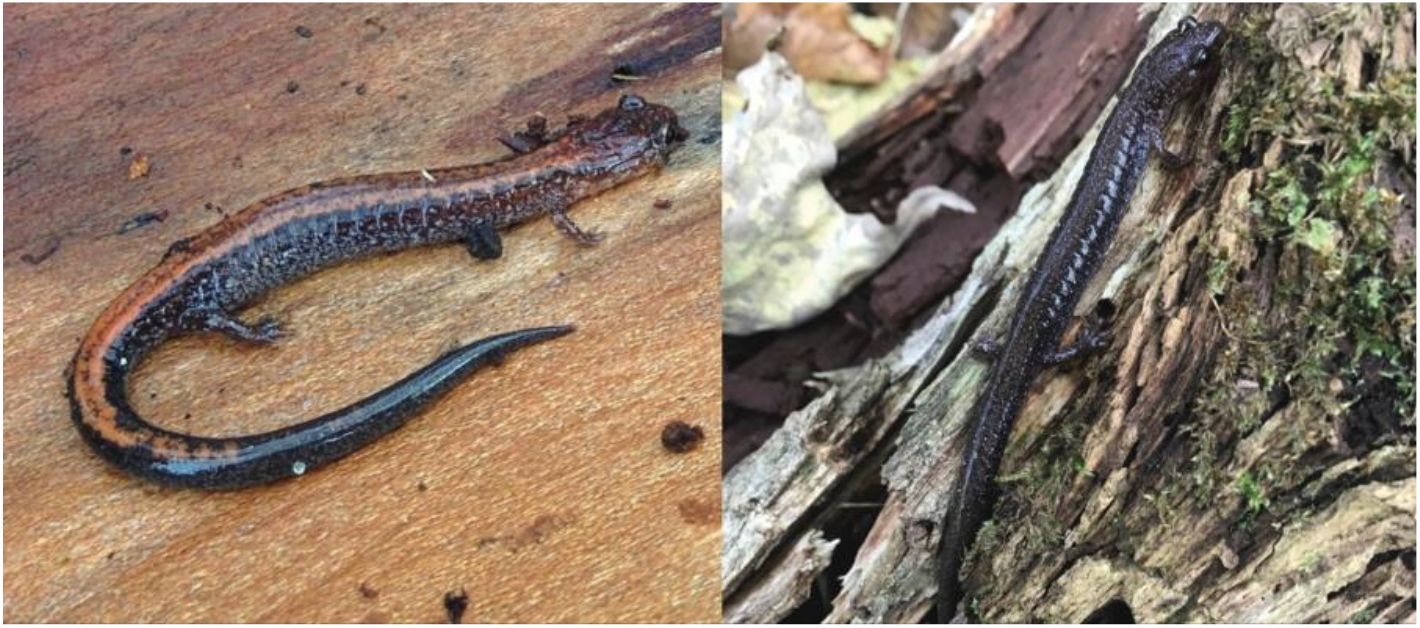
Color morphs of *Plethodon cinereus*. Representative iNaturalist images of the striped (left) and unstriped (right) color morphs of *Plethodon cinereus*. Photos and observations by iNaturalist users Jessica (iNaturalist user jessicapfund) and Myvanwy (iNaturalist user acuriousmagpie), respectively.

The goal of our study was to test range-wide color morph and climate associations by leveraging more than 20,000 community science photographs. We created a computer vision model for scoring striped and unstriped color morphs of *P. cinereus* via an experimental design capable of handling photographs that are highly heterogeneous and vary extensively in quality. With the classified data, we then used ecological niche modeling and a logistic modeling framework to examine whether the two color morphs partition available niche space, thereby contributing to the maintenance of this polymorphism. Our methodological approach not only provides new insight into the association between climate and color morph frequency in *P. cinereus* at the range-wide scale, but also demonstrates a pipeline for rapidly classifying discrete color morphs in community science images. We also discuss the complications faced when developing the computer vision model, but highlight the utility of this approach with continuously growing community science image resources.

## Methods

### Community Science Image Dataset

We downloaded 15,777 research-grade (georeferenced observations with species ID verified by a minimum of two separate reviewers) images of *P. cinereus* from iNaturalist (accessed August 5, 2020) via a command-line query tool (https://gitlab.com/stuckyb/cbg_phenology). Images were not modified in any manner. From this initial set, we randomly selected 4,000 images to be the basis of our training and validation dataset. Seven volunteers aided in scoring salamander color pattern (striped/unstriped). A color pattern scoring guide and training was provided by MMH to all participants prior to scoring to ensure unanimity in trait definitions. Images were divided into 10 sets of 400. All image sets were scored twice by separate volunteers (i.e., no volunteer scored the same image twice). If there was incongruence between volunteers in scoring a color pattern, a third, independent, volunteer provided a consensus score.

To score salamander color patterns, we used the scriptable desktop software program ImageAnt (https://gitlab.com/stuckyb/imageant). We wrote a custom ImageAnt script to query: 1) the number of salamanders in an image; 2) salamander color pattern (striped, unstriped, other); or 3) whether the image was unusable (i.e., the color pattern was unidentifiable). Images with multiple salamanders were subsequently presented with another scoring rubric of “striped”, “unstriped”, or “both color morphs”. In the final training set, images with multiple of the same color morph were lumped with images of a single salamander of the same color morph. Images that contained both color morphs were not included in the training set. Although *P. cinereus* displays a discrete striped/unstriped dorsal color pattern polymorphism, aberrant phenotypes (e.g., leucistic or the orange-red “erythristic’ phenotype) can be found (Moore and Ouellet, 2014). The few cases of erythristic (“other”) phenotypes were included within the “striped” class, while no leucistic examples were observed in our training set. Our final model was trained using the binary categories: “striped” and “unstriped”.

### Deep Learning

We trained a convolutional neural network (CNN) using the EfficientNet (efficientnet-b4; Tan and Le, 2019) architecture implemented in PyTorch with PyTorch Lightning used to implement model training (Falcon, 2019). We implemented transfer learning (Yosinski et al., 2014) with model weights that were pre-trained on the ImageNet dataset (Deng et al., 2009). CNN training and validation was performed on the University of Florida HiPerGator high-performance computer using one GPU.

A series of model training hyperparameters were included and systematically modified to increase validation accuracy. To train the model, we used stochastic gradient descent with momentum and a dynamic learning rate scheduler starting with a learning rate of 0.001 and set to decay by a factor of 0.1 based on validation loss. An oversampling procedure was implemented due to unequal image representation of the striped and unstriped salamander phenotypes. Image preprocessing included resizing images to 596×447 pixels and normalizing the color channels with the same transformation used for ImageNet pretraining. A set of data augmentation techniques was applied to each batch during model training including: 1) random horizontal flips, 2) random vertical flips, 3) random rotations, 4) color jittering, and 5) random affine transforms.

We used k-fold cross-validation with 4 random splits to evaluate model performance. For our final production model, we took the best model from each cross-validation fold (as defined by the lowest validation loss for that fold) and combined them into an ensemble model by averaging the predictions of all four models. Using ImageAnt, we manually scored 500 more images that were independent of those used for model training and validation to serve as a test set for evaluating the final ensemble model. We then used the production ensemble model to analyze all remaining *P. cinereus* images on iNaturalist. Due to the growth of *P. cinereus* research-grade images between model training and validation steps, we re-downloaded all research-grade images from iNaturalist (20,318 images; accessed March 24, 2021) and then analyzed all images not included in the training and test sets using the full model ensemble. Full modeling details and code can be found on our GitHub repository (https://github.com/mhantak/Salamander_image_analysis).

### Environmental Data

To test climatic niche differences between the color morphs of *P. cinereus*, we first obtained bioclimatic (n =19) and elevational data at 30 arc-second (∼1 km) resolution (WorldClim V1.4; Hijmans et al., 2005). We next determined the accessible area for *P. cinereus* by buffering the known geographic range by 100 km and then clipped environmental data layers to that area. After doing so, and to avoid overparameterization and multicollinearity, the environmental data layers were reduced to include only uncorrelated variables (r = .80). The final dataset included eight variables: elevation, mean diurnal range (BIO2), maximum temperature of warmest month (BIO5), temperature annual range (BIO7), mean temperature of wettest quarter (BIO8), mean temperature of direst quarter (BIO9), precipitation seasonality (BIO15), and precipitation of warmest quarter (BIO18).

### Niche Modeling

We used ecological niche modeling (ENM) as a means to determine niche characteristics of both morphs. Prior to running niche models, we first filtered the iNaturalist data records. Filtering included removing records with missing or incomplete latitude and longitude information, duplicate records, and manually removing records outside of the known range. To reduce the potential for spatial autocorrelation and bias from areas with particularly dense sampling, we thinned our data to include records separated by a minimum of 25 kilometers. ENM’s were constructed separately for both the striped and unstriped morphs using the maximum entropy algorithm implemented in MAXENT V3.4.1 (Phillips et al., 2006) in the R package *ENMeval* (Muscarella et al., 2014). Data were partitioned using the “block” method to account for spatial autocorrelation. Regularization multipliers ranged from 0.5 to 5 and possible feature combinations were: L, H, LP, LQ, LQH, LQP, and LQPH (L = linear, H = hinge, P = product, Q = quadratic). The best model was selected based on the lowest ΔAICc. After model calibration and validation, we converted the modeled output of predicted probabilities of presence within the accessible area to binary presence/absence maps using equate entropy and the original distribution (cloglog) threshold, which typically performs well when attempting to balance omission error versus the fraction of predicted presence. We next examined niche overlap between the two morphs with the Schoener’s *D* metric using *ENMeval*. Niche breadth of both morphs was calculated using the raster.breadth function in the R package ENMTools (Warren et al., 2010). To visually examine color morph overlap in association with climatic predictors, we ran a principal component analysis (PCA) using the reduced set of bioclimatic variables and the predicted presence points from the striped and unstriped morph ENM’s with the base R prcomp() function (R Core Team, 2019).

### Statistical Analyses

We further quantified niche differences between the morphs by running a multiple logistic regression using the R base glm() function (R Core Team, 2019) with a binomial family and a logit link function. The predictors for this model were generated by assembling underlying bioclimatic conditions (e.g., BIO2, BIO5, BIO7, BIO8, BIO9, BIO15, BIO18) and elevation at each pixel predicted as a presence in the above binarized maps, for both morphs. We opted to use the raw environmental data rather than principal components for ease of interpretation. Color morph, coded as 1 for striped morphs and 0 for unstriped morphs was the response variable. All predictors were mean-centered and scaled. In order to select the best model, and given no *a priori* hypotheses about the best predictors, we used the ‘dredge’ function in the R package *MuMIn* (Barton, 2012) to rank and assess the best-fit model with AICc. If any predictors were not in the top model or if any predictor variance inflation factor (VIF) was greater than four, we dropped those variables and re-ran the logistic regression. To generate a pseudo-R^2^ value, as a measure of goodness of fit for our best-fit model, we used the ‘r2_nagelkerke’ function in the R package *performance* (Lüdecke et al., 2021).

## Results

### Volunteer and Model Accuracy

Across the seven volunteers that scored the 4,000 training and validation images, we estimate that mean volunteer annotation accuracy was 95.9%. Consensus was achieved for 3,871 (3,005 striped, 866 unstriped) images, while the remaining 129 images were either unidentifiable as striped or unstriped salamanders (n=51) or unusable because both morphs were visible in the image (n=78; Table 1). The majority of images were scored with a mean scoring time of three seconds. Some images took annotators considerably longer to analyze, although extremely long annotation times were likely due to annotators leaving ImageAnt running while not actively scoring. The 3,871 images served as the basis for model training and validation.

**Table 1.**
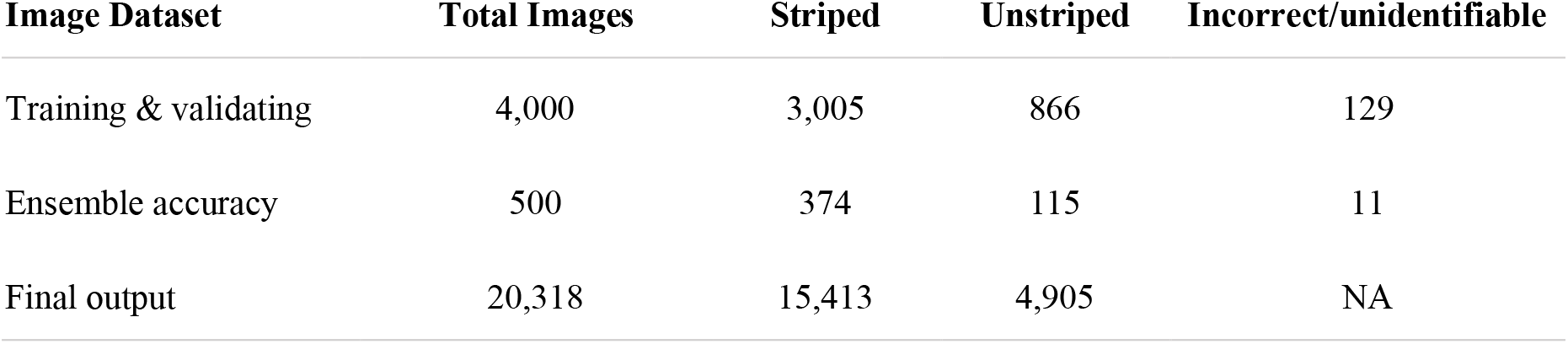
Total number of images and how the images were classified for different datasets. 1) Volunteer scoring for the training and validation dataset. 2) Examination of a subset (500 images) of the final model ensemble to retrieve an estimate of model accuracy. 3) Final computer vision model color pattern scores.

| Image Dataset | Total Images | Striped | Unstriped | Incorrect/unidentifiable |
| --- | --- | --- | --- | --- |
| Training & validating | 4,000 | 3,005 | 866 | 129 |
| Ensemble accuracy | 500 | 374 | 115 | 11 |
| Final output | 20,318 | 15,413 | 4,905 | NA |

Validation accuracy across the four cross-validation folds varied minimally (fold 1 = 98.6%; fold 2 = 97.3%; fold 3 = 96.2%; fold 4 = 97.4%). The mean cross-validation accuracy was 97.4% and the test accuracy of the final ensemble model was 97.8% (Table 1). Out of the 20,318 iNaturalist images analyzed by the ensemble model, 15,413 (75.9%) were labeled as striped and 4,905 (24.1%) as unstriped salamanders (Table 1)

### Niche Modeling

Our filtering steps removed 60 data points, generating a final dataset of 20,258 total point presences (N = 15,363 striped morphs; N = 4,895 unstriped morphs; Fig. 2). These were used along with the uncorrelated environmental predictors to generate a best-fit MAXENT model for striped and unstriped morphs. The best model for both striped and unstriped, based on AICc and ΔAICc, consisted of LQPH features with a regularization multiplier of two (striped model AICc =28877.63, ΔAICc = 4.86; unstriped model AICc = 16072.40, ΔAICc = 6.09). AUC_train_ (striped 0.78; unstriped 0.82) suggests relatively performant models; because *P. cinereus* is widespread and common across its range, separating higher and lower quality habitat is more challenging than for habitat specialists. AUC_test_ values (striped 0.75; unstriped 0.81) were close to the AUC_train_ scores, suggesting these models are not overfit. The Schoener’s *D* metric indicates that the niches of the morphs overlap at 87%. Niche breadth of the striped morph is greater than that of the unstriped morph (Levins B2; striped = 0.64; unstriped = 0.55). The PCA of the reduced bioclimatic variables shows how the morphs partition niche space (Table S1, Fig. 3). PC1 represents 30% of the variation and its loadings are primarily mean diurnal range (BIO2), maximum temperature of warmest month (BIO5), and precipitation of warmest quarter (BIO18; Fig 3A). PC2 represents 26% of the variation and mean temperature of driest quarter (BIO9), temperature annual range (BIO7), and precipitation seasonality (BIO15; Fig. 3A) are the main loadings. Lastly, 17% of the variation is explained by PC3, with loadings primarily from elevation (ALT) and maximum temperature of warmest month (BIO5; Fig. 3B).

**Figure 2.**
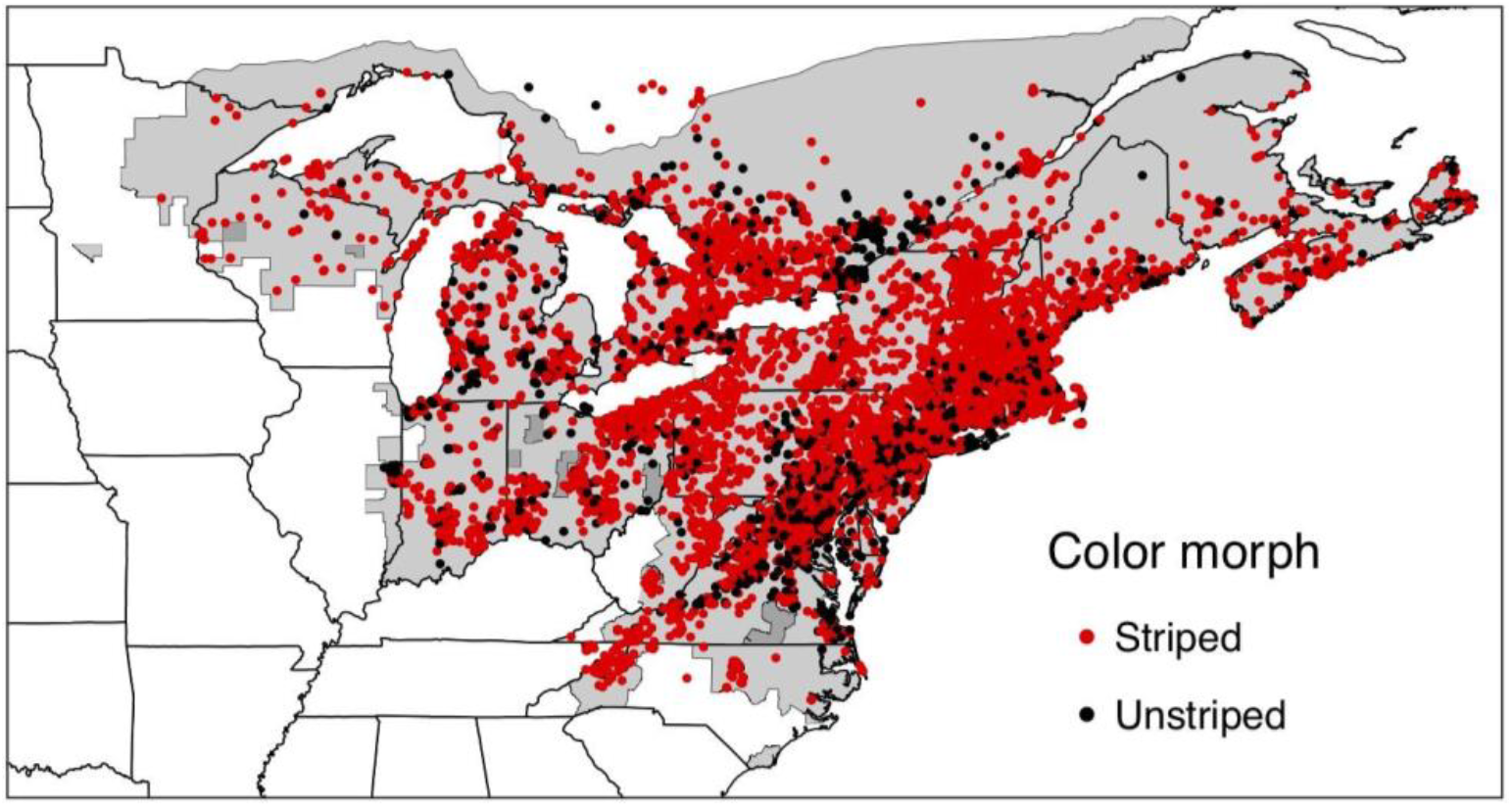
Color morph data generated from the computer vision model. Georeferenced iNaturalist observations (N = 20,258) of *P. cinereus*. Record localities are colored by morph (red = striped, black = unstriped) based on the final computer vision model run.

**Figure 3.**
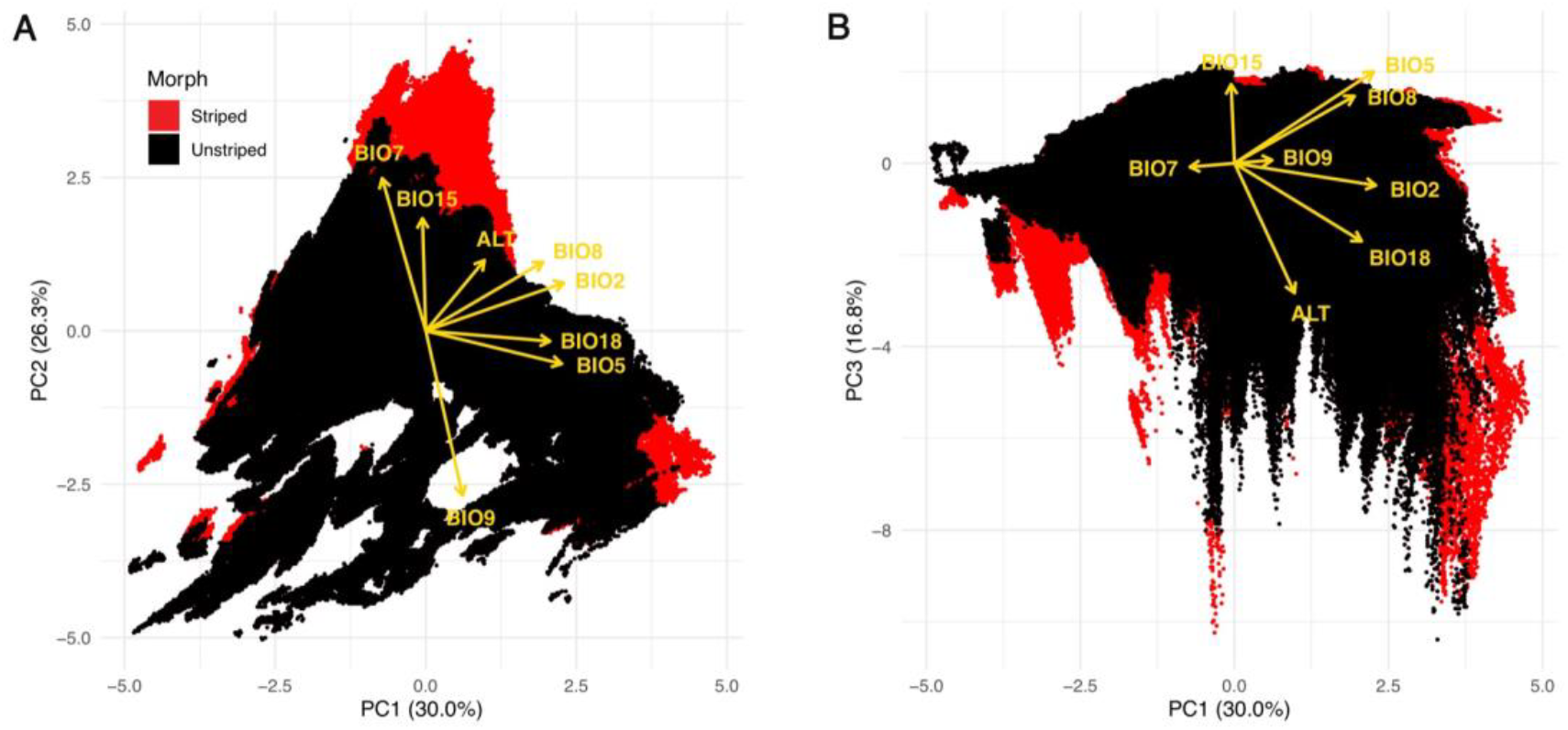
Climatic niche differences between color morphs of *Plethodon cinereus*. PCA of reduced climatic variables: A) PC1-PC2, B) PC1-PC3. Predicted presence points from striped and unstriped morph ecological niche models were grouped into hexbins (red = striped; black = unstriped). PCA loadings are represented by yellow arrows.

### Logistic Modeling

The best model included elevation and all seven bioclimatic predictors (BIO2, BIO5, BIO7, BIO8, BIO9, BIO15, BIO18), however, BIO5 was subsequently dropped because it had a VIF greater than four (PsuedoR^2^ = 0.04). All model effects were significant. Striped morph frequency is positively correlated with elevation (*β* = 0.051, *SE* = 0.001, *p* < 0.001; Fig. 4A). There is a decreased odds of striped morphs with mean diurnal range (BIO2; *β* = -0.063, *SE* = 0.001, *p* < 0.001; Fig. 4B). Striped morph frequency has higher odds of occurring with higher temperature annual range (BIO7; *β* = 0.126, *SE* = 0.001, *p* < 0.001; Fig. 4C), but the odds decrease with mean temperature of the wettest quarter (BIO8; *β* = -0.040, *SE* = 0.001, *p* < 0.001; Fig. 4D). The odds of striped morph frequency increases with mean temperature of driest quarter (BIO9; *β* = 0.112, *SE* = 0.001, *p* < 0.001; Fig. 4E) and with both precipitation predictors: precipitation seasonality (BIO15; *β* = 0.315, *SE* = 0.001, *p* < 0.001; Fig. 4F) and precipitation of the warmest quarter (BIO18; *β* = 0.178, *SE* = 0.001, *p* < 0.001; Fig. 4G). Precipitation effect sizes were generally stronger than temperature in separating morphs.

**Figure 4.**
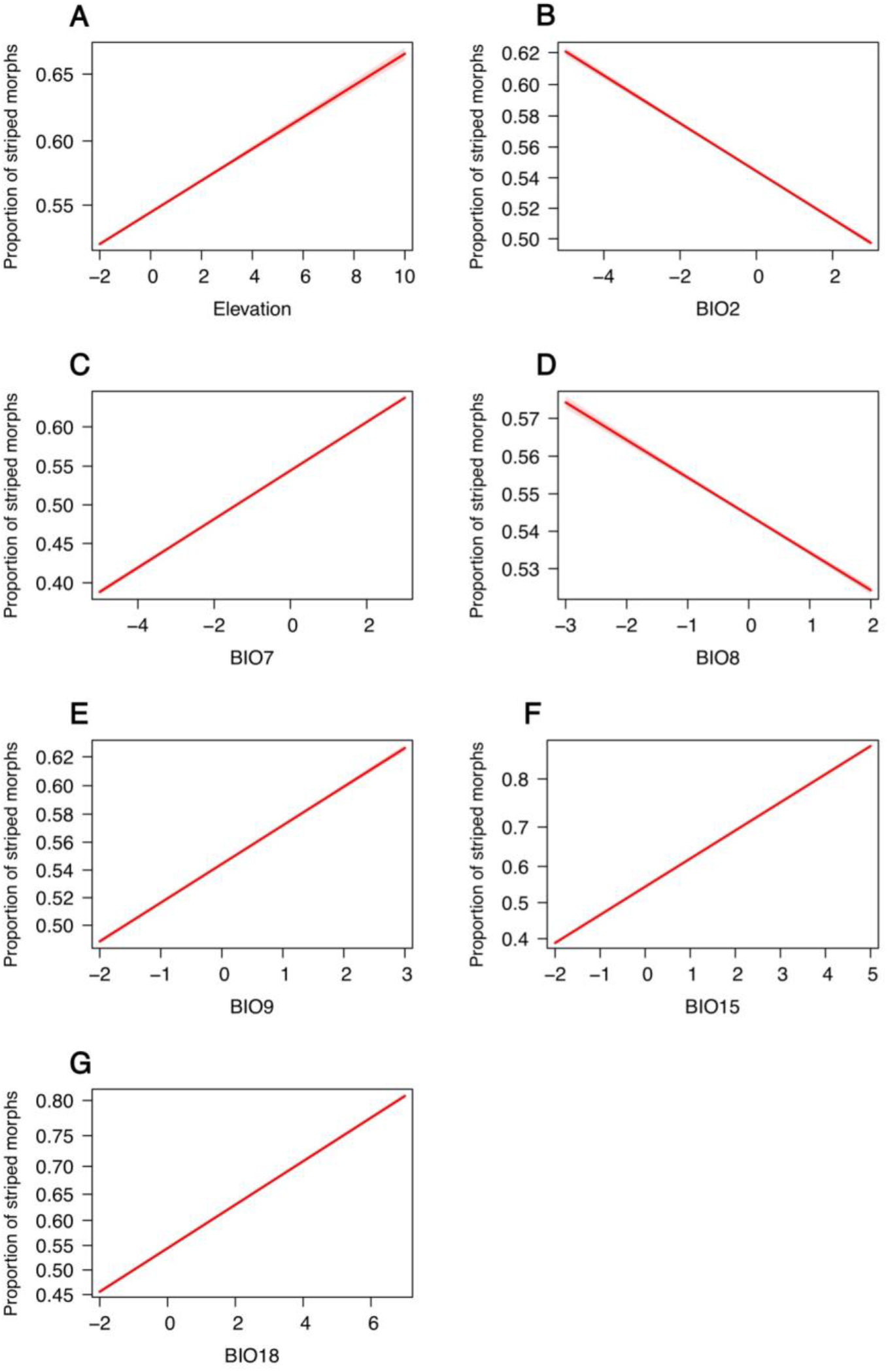
Climatic predictors of color morph frequency. Top model effect plots of color morph frequency variation in *P. cinereus*. The proportion of color morphs is influenced by A) elevation; B) mean diurnal range (BIO2); C) temperature annual range (BIO7); D) mean temperature of wettest quarter (BIO8); E) mean temperature of direst quarter (BIO9); F) precipitation seasonality (BIO15); and G) precipitation of warmest quarter (BIO18). 95% confidence intervals are included in each plot.

## Discussion

Community science resources, especially images tied to community identifications available via iNaturalist, are rapidly expanding. These images contain a treasure trove of biologically relevant information about phenotypes and interactions (DiCecco et al., 2021), but unlocking this information remains a challenge. Thus far, computer vision models have largely focused on species identification from images (Gomez Villa et al., 2017; Norouzzadeh et al., 2018; Willi et al., 2019). To our knowledge, no previous studies have aimed to use machine learning approaches to extract trait information, but such approaches are needed given the deluge of records with digital vouchers being submitted. Here, we created a highly accurate (∼98% accurate based on test set evaluation) computer vision model for classifying a salamander’s color pattern from community science images. With the data produced from this model, we expanded our knowledge of why a common striped/unstriped color polymorphism persists in the abundant salamander, *Plethodon cinereus*.

### Scalability of Community Science Images

A challenge of using CNNs for feature classification is the need for robust sample sizes for training. Community science platforms, such as iNaturalist, hold millions of images of various plants and animals that are spatially and temporally replicated. A well-established machine learning algorithm provides iNaturalist users with a suggested species identification (www.inaturalist.org/). A few studies have manually scored traits such as flower presence or absence in order to identify phenological patterns across geography (Barve et al., 2020; Li et al., 2021). Yet, manual scoring of more images would be necessary to expand upon these studies. Our pipeline provides a streamlined example of how to obtain large-scale trait data from community science images. This computer vision model can now be used to rapidly score the trait of interest, and can be used in perpetuity to gather data on more records as they become available on community science platforms. From August 5th, 2020 when we downloaded our core image dataset used for model training to January 13th, 2022, the number of research-grade *P. cinereus* records has nearly doubled (from 15,777 to 29,040). As well, many other *Plethodon* species have similar color polymorphisms and our model should be transferable to these other species.

Community science images are not perfect. With unstandardized images, expert decisions on feature classifications are key. For this work, we created a salamander color scoring guide (found in https://github.com/mhantak/Salamander_image_analysis) that was distributed to all volunteers who aided in creating the training dataset. While standardization of training data is important, some aspects of community science images remain out of our control and create unique challenges when designing machine learning experiments. For instance, during volunteer scoring, there were a few research-grade species misidentifications, which is unsurprising given that closely related species can look nearly identical to *P. cinereus* (Fisher-Reid and Wiens, 2015). These sorts of issues are inherent in working with community science data, but careful consideration is needed when making decisions about how to deal with these records. In our case, we scored misidentified species to the most similar looking morph of *P. cinereus*. For example, a Two-lined Salamander (*Eurycea* sp.), was categorized as a striped morph, while a Slimy Salamander (*Plethodon glutinosus*) was scored as an unstriped morph. Keeping these images of similar-looking species in the training dataset provides a more representative sample of what the model will encounter when analyzing new images. Further, there were several images solely of the ventral side of the salamander. While not a misidentification, the needed trait information is best obtained from a dorsal view, and ventral views would be better suited as additional images to augment iNaturalist records that also include a dorsal view. Due in part to these ventral images, there were 51 images out of 4,000 (1.3%) that were excluded from the training dataset because they could not be identified to morph. Other image problems included excessive blurriness, partial body part exposure (e.g., head only), or a salamander that was too distant in the photograph. Even if ∼1% of all input images are unidentifiable and the model were to incorrectly guess on all of them, we maintain that this is still an acceptable error rate when dealing with community science images. Finally, we removed one extraneous data point from the data after determining it was well outside of the geographic range of the species. One record out of >20,000 is a very low error rate.

### Computer Vision Model Intricacies

Our final computer vision model is based on a binary classification, ‘striped’ or ‘unstriped’ color morph. This simplified binary classifier works for the majority of individual *P. cinereus* across the distribution of the species. However, there is a third, uncommon erythristic (orange-red) color morph, which we combined with the striped morph (similar to another study; Fisher-Reid and Wiens, 2015) because there were too few examples in our training image set (n = 20) to train a model to identify it. In addition, other abnormal color phenotypes of *P. cinereus* can sometimes be found (see Moore and Ouellet, 2014). When preparing our training dataset, we found 16 instances of a white (instead of orange or red) striped phenotype. As with the erythristic phenotype, these images were too sparse for model training and were lumped with striped morphs based on the existence of the dorsal stripe. Similar decisions were necessary for less frequent aberrant phenotypes. Single images that contained multiple salamanders also posed an issue with creating our training set. We initially considered attempting to train a model to determine the number of salamanders in an image or identify images with multiple salamanders. However, a stepwise classifier would require more training images for the additional categories and ultimately create a more unbalanced dataset, as there were less images with multiple salamanders. We, thus, adopted the simple solution of combining images with multiple salamanders of the same phenotype with images of single salamanders (e.g., an image with three striped morphs was binned into the “striped” class). We removed images that contained both color morphs from the training set because *either* category (striped or unstriped) could be considered correct for these images. At inference time, images with both color morphs were considered to be correctly classified regardless of which color morph the model assigned them. Such images are quite rare and accounted for only 78 of the 4,000 images analyzed to generate the training and test sets.

### Climate & Color Morph Trends in the Eastern Red-backed Salamander

The ecological niche models show that the morphs largely overlap (i.e., by 87%) in climatic niche space, but striped morphs have a wider niche breadth than unstriped morphs. The PCA highlights the variation between *P. cinereus* color morphs and in general shows that striped morphs can be found in areas with more variable climatic conditions. Logistic model findings are consistent with the PCA and demonstrate a positive association between striped morph frequency and elevation, metrics of precipitation, and two climate variables (BIO7 and BIO9). Whereas the proportion of striped morphs decreases with mean diurnal range (BIO2) and mean temperature of wettest quarter (BIO8).

Our finding of a positive relationship between elevation and striped morph frequency is consistent with previous studies (Gibbs and Karraker, 2006; Moore and Ouellet, 2015; Hantak et al., 2021). Following the expectation that higher elevations are typically colder than lower elevations, we predicted the observed positive correlation. However, here and in other studies, striped morphs are not always associated with cooler temperatures. A recent study by Hantak et al. (2021) found the proportion of striped morphs increases with increasing elevation and mean annual temperature and, based on these results suggested that these predictors may be decoupled in relation to color morph frequency in *P. cinereus*. While the reason for greater proportion of striped morphs in higher elevations remains unclear, it may be possible that gene flow is reduced along altitudinal gradients in this species. Previous work has shown that elevation is a significant predictor of genetic differentiation in amphibians (Funk et al., 2005; Giordano et al., 2007; García-Rodríguez et al., 2021), including *P. cinereus* (Hantak et al., 2019); although moderate changes in elevation was not the most important driver of morph frequency variation in northern Ohio (Hantak et al., 2019).

Based on previous studies of climate associations between in *P. cinereus* color morphs, we predicted that striped morph occurrences would be more tightly linked with cooler and wetter climatic niches, whereas unstriped morphs would be more correlated with warmer, drier niches (Lotter and Scott, 1977; Moreno, 1989; Anthony et al., 2008). While we found the predicted trend for precipitation with striped morph frequency, our temperature-morph findings were more nuanced. The PCA and logistic model indicates that the striped morph is, in general, found in areas with more variability in temperature. Whereas the proportion of striped morphs decreases with mean diurnal range (BIO2), suggesting that striped morphs are negatively impacted by temperature fluctuations. In addition, the proportion of striped morphs decreased with mean temperature of wettest quarter (BIO8), indicating a possible humidity threshold for this morph.

Much work on the polymorphism in *P. cinereus* relies heavily on findings that were conducted over relatively small spatial and temporal scales. In addition, some studies have found no climate-related morph trends or inconsistent patterns over time (Petruzzi et al., 2006; Muñoz et al., 2016; Evans et al., 2018). Fisher-Reid et al. (2013) demonstrated that striped morphs were found in warmer, wetter habitats on Long Island, New York, while Hantak et al. (2021) found striped morphs were more associated with warmer, drier habitats in localities across Maryland, New York, and Virginia. Range-wide, dense data can help examine overall trends and localize those at finer scale in a unifying framework. Besides our current work, two other studies have attempted to examine climate-morph trends in *P. cinereus* across a greater proportion of the species range. But here again, these studies find conflicting results likely due to differences in datasets, covariates, and statistical approaches (Gibbs and Karraker, 2006; Moore and Ouellet, 2015; Cosentino et al., 2017). It is possible that these variations in approaches lead to ambiguous color morph and climate relationships, or it may be there are more complex contextual cues that are being missed when attempting to understand polymorphism rates in *P. cinereus*. With physiological differences between the morphs (Moreno, 1989; Davis and Milanovich, 2010; Smith et al., 2015), climate likely plays some role in morph distribution, but other, local, selection pressures may be more important in this system.

### Next Steps

The combination of community science and deep learning provides a powerful resource for future studies of phenotypic variation. With the rapid growth of data, including community science images, scalable resources such as computer vision models are necessary to keep pace with rate of data accumulation (Hassoun et al., 2021), which potentially provides a means to track temporal changes, not simply spatial ones. A further step for our research is to use this model to score color morphs of other species within the salamander genus *Plethodon*. In total, there are 10 species within *Plethodon* that contain the exact same dorsal striped/unstriped color pattern (Petranka, 1998; Highton, 2004). Occurrence data points are available for all of these other species on the iNaturalist platform, ranging from ∼70 observations for the IUCN listed “vulnerable” mountaintop endemic, *P. sherando* (Highton and Collins, 2006) to >2,000 observations of the more widespread Western Red-backed Salamander (*P. vehiculum*). Much research has been done on the morphs of *P. cinereus*, but very little is known about dynamics of the polymorphism in these other species, including whether the morphs diverge in climatic niche space. Although our computer vision model was developed to score salamander striped and unstriped color patterns, our entire workflow can also be transferred to any system that has discrete, easily identifiable, trait variation.

### Limitations of the Study

Although machine learning holds much promise for rapidly gathering phenotypic data from digital images, the main limitation to using fully supervised deep learning approaches is the number of labeled training images (and, primarily, the time and expertise needed to generate the labels). Depending on the complexity of the intended classification, several thousand vouchers for each category may be necessary for training and validating the model. Here, we present a relatively simple problem: are the salamanders striped or unstriped? Adding categories or addressing more complicated phenotypes will require more training images. In the deep learning literature, methods to reduce the labeling bottleneck (e.g., through one- or few-shot learning; O’Mahony et al. 2019; Wang et al. 2020) are being developed and future studies on the applicability and effectiveness of those methods to the application presented here are needed. The other main limitation to the type of work we presented in this paper is the imperfect nature of community science images. Misidentifications do occur, even when reducing the dataset to vetted (e.g., research-grade) images, and images themselves vary in absolute quality and relative usability for a particular trait scoring outcome. Solutions to dealing with these issues will be on a case-to-case basis, but in our work, we found that labeling misidentified species to the closest phenotype and filtering some of the most problematic images worked well. Misidentifications and unusable images are inherent when working with community science data, but they are infrequent. With tens of thousands of correctly identified images of usable quality, a few misidentifications and image issues will not dramatically impact the biological conclusions of the study. Certainly, future work can also include leveraging weak-learning approaches that are more robust to the presence of label noise and inaccuracies.

## Resource Availability

### Lead Contact

Further information or requests for resources should be directed to the Lead Contact, Maggie M. Hantak

## Data and Code Availability

Data and code for this study are available at https://github.com/mhantak/Salamander_image_analysis.

## Acknowledgements

We thank iNaturalist community science users for their efforts in documenting and identifying Eastern Red-backed Salamander occurrences. We also thank volunteers N. Barve, V. Barve, N. Gardner, and N. Sewnath for scoring salamander color patterns, which were used in the training dataset. Research reported in this publication was supported by the University of Florida Informatics Institute Fellowship Program and the NSF Postdoctoral Research Fellowship in Biology (2010776) both awarded to MMH.

## Author Contributions

MMH, RPG, AZ, and BJS designed the study. MMH and BJS developed the computer vision model. Ecological analyses were led by MMH and RPG. MMH wrote the manuscript. All authors contributed to drafts and the final version of the manuscript.

## Supplemental Information

**Supplemental Table 1.** PCA bioclimatic variable loadings from the reduced set of elevation (ALT), temperature (BIO1-BIO9, and precipitation (BIO15 and BIO18) covariates. The standard deviation, proportion of variance, and cumulative proportion is provided for each PC axis.

| Variable | PC1 | PC2 | PC3 | PC4 | PC5 | PC6 | PC7 | PC8 |
| --- | --- | --- | --- | --- | --- | --- | --- | --- |
| ALT | 0.217 | 0.256 | -0.629 | 0.352 | -0.070 | 0.476 | 0.101 | 0.356 |
| BIO2 | 0.510 | 0.174 | -0.107 | -0.499 | 0.197 | 0.336 | 0.011 | -0.545 |
| BIO5 | 0.502 | -0.119 | 0.444 | -0.166 | -0.051 | 0.181 | -0.369 | 0.581 |
| BIO7 | -0.161 | 0.552 | -0.018 | -0.517 | 0.190 | -0.236 | 0.351 | 0.432 |
| BIO8 | 0.432 | 0.249 | 0.326 | 0.292 | -0.469 | -0.201 | 0.531 | -0.130 |
| BIO9 | 0.136 | -0.596 | 0.015 | 0.003 | 0.447 | 0.089 | 0.626 | 0.160 |
| BIO15 | -0.012 | 0.407 | 0.383 | 0.482 | 0.653 | 0.117 | -0.081 | -0.091 |
| BIO18 | 0.458 | -0.037 | -0.378 | 0.125 | 0.265 | -0.714 | -0.221 | 0.027 |
| Standard deviation | 1.548 | 1.450 | 1.161 | 0.872 | 0.793 | 0.680 | 0.439 | 0.330 |
| Proportion of variance | 0.300 | 0.263 | 0.168 | 0.095 | 0.079 | 0.058 | 0.024 | 0.014 |
| Cumulative proportion | 0.300 | 0.563 | 0.731 | 0.826 | 0.905 | 0.962 | 0.986 | 1.000 |

## References

Anthony, C. D., Venesky, M. D., and Hickerson, C. A. M. (2008). Ecological separation in a polymorphic terrestrial salamander. J. Anim. Ecol. 77, 646–653.

Barton, K. (2012). Package ‘MuMIn’. Model selection and model averaging based on information criteria. R package version 3.2.4. http://cran.r-project.org/web/packages/MuMIn/index.html.

Barve, V. V., Brenskelle, L., Li, D., Stucky, B. J., Barve, N. V., Hantak, M. M., McLean, B. S., Paluh, D. J., Oswald, J. A., Belitz, M. W., Folk, R. A., and Guralnick, R. P. (2020). Methods for broad-scale plant phenology assessments using citizen scientists’ photographs. Appl. Plant Sci. 8, e11315.

Cosentino, B. J., Moore, J.-D., Karraker, N. E., Ouellet, M., and Gibbs, J. P. (2017). Evolutionary response to global change: climate and land use interact to shape color polymorphism in a woodland salamander. Ecol. Evol. 7, 5426–5434.

Davis, A. K., and Milanovich, J. R. (2010). Lead-phase and red-stripe color morphs of red-backed salamanders Plethodon cinereus differ in hematological stress indices: a consequence of differential predation pressure? Curr. Zool. 56, 238–243.

Deng, J., Dong, W., Socher, R., Li, L. J., Li, K., and Fei-Fei, L. (2009). ImageNet: A Large-Scale Hierarchical Image Database. In 2009 IEEE Conference on Computer Vision and Pattern Recognition, 248–255.

DiCecco, G. J., Barve, V., Belitz, M. W., Stucky, B. J., Guralnick, R. P., and Hurlbert, A. H. (2021). Observing the observers: How participants contribute data to iNaturalist and implications for biodiversity science. Bioscience 71, 1179–1188.

Endler, J. A., and Mappes, J. (2017). The current and future state of animal coloration research. Philos. Trans. R. Soc. Lond. B. Biol. Sci. 372, 20160352.

Evans, A. E., Forester, B. R., Jockusch, E. L., and Urban, M. C. (2018). Salamander morph frequencies do not evolve as predicted in response to 40 years of climate change. Ecography 41, 1687–1697.

Falcon, W. (2019). Pytorch lightning. GitHub. Note: https://github.com/williamFalcon/pytorch-lightning.

Fisher-Reid, M. C., and Wiens, J. J. (2015). Is geographic variation within species related to macroevolutionary patterns between species? J. Evol. Biol. 28, 1502–1515.

Fisher-Reid, M. C., Engstrom, T. N., Kuczynski, C. A., Stephens, P. R., and Wiens, J. J. (2013). Parapatric divergence of sympatric morphs in a salamander: incipient speciation on Long Island? Mol. Ecol. 22, 4681–4694.

Ford, E. B. (1945). Polymorphism. Biol. Rev. 20, 73–88.

Funk, W. C., Blouin, M. S., Corn, P. S., Maxell, B. A., and Pilliod, D. S. Population structure of Columbia spotted frogs (Rana lueteiventris) is strongly affected by the landscape. (2005). Mol. Ecol. 14, 1–14.

García-Rodríguez, A., Martínez, P. A., Oliveira, B. F., Velasco, J.A., Pyron, R. A., and Costa, G. C. (2021). Amphibian speciation rates support a general role of mountains as biodiversity pumps. Am Nat. 198, E68–E79.

Gibbs, J. P., and Karraker, N. E. (2006). Effects of warming conditions in east-ern North American forests on red-backed salamander morphology. Conserv. Biol., 20, 913–917.

Giordano, A.R., Ridenhour, B. J., and Storfer, A. (2007). The influence of altitude and topography on genetic structure in the long-toed salamander (Ambystoma macrodactulym). Mol. Ecol. 16, 1625–1637.

Gomez Villa, A., Salazar, A., and Vargas, F. (2017). Towards automatic wild animal monitoring: Identification of animal species in camera-trap images using very deep convolutional neural networks. Ecol. Inform. 41, 24–32.

Hantak, M. M., Page, R. B., Converse, P. E., Anthony, C. D., Hickerson, C. M., and Kuchta, S. R.. (2019). Do genetic structure and landscape heterogeneity impact color morph frequency in a polymorphic salamander? Ecography 42, 1383–1394.

Hantak, M. M., Federico, N. A., Blackburn, D. C., and Guralnick, R. P. (2021). Rapid phenotypic change in a polymorphic salamander over 43 years. Sci. Rep. 11, 22681.

Hassoun, A., Jefferson, F., Shi, X., Stucky, B., Wang, J., and Rosa Jr., E. (2021). Artificial intelligence for biology. Integrative and Comparative Biology doi.org/10.1093/icb/icab188.

Highton, R. (2004). A new species of woodland salamander of the Plethodon cinereus group from the Blue Ridge Mountains of Virginia. Jeffersoniana 14, 1–22.

Highton, R., and Collins, J. (2006). Plethodon sherando. The IUCN Red List of Threatened Species 2006: e.T61905A12569864.

Hijmans, R. J., Cameron, S. E., Parra, J. L., Jones, P. G., and Jarvis, A. (2005). Very high resolution interpolated climate surfaces for global land areas. Int. J. Climatol. 25, 1965–1978.

iNaturalist. (2021). Available online: https://www.inaturalist.org/.

Lattanzio, M. S., and Buontempo, M. J. (2021). Ecogeographic divergence linked to dorsal coloration in Eastern Hog-Nosed Snakes (Heterodon platirhinos). Herpetologica 77, 134–145.

LeCun, Y., Bengio, Y., and Hinton, G. (2015). Deep learning. Nature 521, 436–444.

Lehtinen, R. M., Carlson, B. M., Hamm, A. R., Riley, A. G., Mullin, M. M., and Gray, W. J. (2020). Dispatches from the neighborhood watch: Using citizen science and field survey data to document color morph frequency in space and time. Ecol. Evol. 10, 1526.

Li, D., Barve, N., Brenskelle, L., Earl, K., Barve, V., Belitz, M. W., Doby, J., Hantak, M. M., Oswald, J. A., Stucky, B. J., Walters, M., and Guralnick, R. P. (2021). Climate, urbanization, and species traits interactively drive flowering duration. Glob. Change Biol. 27, 892–903.

Lotter, F., and N. J. Scott Jr. (1977). Correlation between climate and distribution of the color morphs of the salamander Plethodon cinereus. Copeia 1977, 681–690.

Lüdecke, D., Ben-Shachar, M., Patil, I., Waggoner, P., and Makowski, D. (2021). performance: An R Package for Assessment, Comparison and Testing of Statistical Models. J. Open Source Softw. 6, 3139.

Lürig, M., Donoughe, S., Svensson, E. I., Porto, A., and Tsuboi, M. (2021). Computer vision, machine learning, and the promise of phenomics in ecology and evolutionary biology. Front. Ecol. Evol. 9, 642774.

Mckinnon, J. S., and Pierotti, M. R. (2010). Colour polymorphism and correlated characters: genetic mechanisms and evolution. Mol. Ecol. 19, 5101–5125.

McLean, C. A., and Stuart-Fox, D. (2014). Geographic variation in animal colour polymorphisms and its role in speciation. Biol. Rev. 89, 860–873.

Moore, J. D., and Ouellet, M. (2014). A review of colour phenotypes of the eastern red-backed salamander, Plethodon cinereus, in North America. Can. Field-Nat. 128, 250–259.

Moore, J. D., and Ouellet, M. (2015). Questioning the use of an amphibian colour morph as an indicator of climate change. Glob. Change Biol. 21 566–571.

Moreno, G. (1989). Behavioral and physiological differentiation between the color morphs of the salamander, Plethodon cinereus. J. Herpetol. 23, 335–341.

Muñoz, D. J., Hesed, K. M., Grant, E. H. C., and Miller, D. A. W. (2016). Evaluating within-population variability in behavior and demography for the adaptive potential of a dispersal-limited species to climate change. Ecol. Evo. 6, 8740–8755.

Muscarella, R., Galante, P. J., Soley-Guardia, M., Boria, R. A., Kass, J. M., Uriarte, M., and Anderson, R. P. (2014). ENMeval: An R package for conducting spatially independent evaluations and estimating optimal model complexity for MAXENT ecological niche models. Methods Ecol. Evol., 5, 1198–1205.

Norouzzadeh, M. S., Nguyen, A., Kosmala, M., Swanson, A., Palmer, M. S., Packer, C., and Clune, J. (2018). Automatically identifying, counting, and describing wild animals in camera-trap images with deep learning. Proc. Natl. Acad. Sci., 115, E5716–E5725.

O’ Mahony, N., Campbell, S., Carvalho, A., Krpalkova, L., Hernandez, G. V., Harapanahalli, S., Riordan, D., and Walsh, J. (2019). One-shot learning for custom identification tasks; A review. In Procedia Manufacturing, Elsevier B.V.: Amsterdam, The Netherlands. Volume 38, pp. 186–193.

Peña, J., Gutiérrez, P., Hervás-Martínez, C., Six, J., Plant, R., and López-Granados, F. (2014). Object-based image classification of summer crops with machine learning methods. Remote Sens. 6, 5019.

Petranka, J. W. 1998. Salamanders of the United States and Canada. Smithsonian Press, Washington.

Petruzzi, E. E., Niewiarowski, P. H., and Moore, F .B.-G. (2006). The role of thermal niche selection in maintenance of colour polymorphism in redback salamanders (Plethodon cinereus). Front. Zool. 3, 10.

Phillips, S. J., Anderson, R. P., and Schapire, R. E. (2006). Maximum entropy modeling of species geographic distributions. Ecol. Modell. 190, 231–259.

Pollicelli, D., Coscarella, M., and Delrieux, C. (2020). RoI detection and segmentation algorithms for marine mammals photo-identification. Ecol. Inform. 56, 101038.

R Core Team. (2019). R: a language and environment for statistical computing. R Foundation for Statistical Computing, https://www.R-project.org/.

Sermanet, P., Frome, A., and Real, E. (2014). Attention for fine grained categorization. 1412.7054.

Sinervo, B., and E. Svensson. (2002). Correlational selection and the evolution of genomic architecture. Heredity 89, 329–338.

Smith, G. R., Johnson, T., Smith, W. O. (2015). Effects of colour morph and season on the dehydration and rehydration rates of Plethodon cinereus. Amphib-Reptil. 36, 170–174.

Tabak, M. A., Norouzzadeh, M. S., Wolfson, D. W., Sweeney, S. J., Vercauteren, K. C., Snow, N. P., Halseth, J. M., Di Salvo, P. A., Lewis, J. S., White, M. D., et al. (2019). Machine learning to classify animal species in camera trap images: Applications in ecology. Methods Ecol. Evol. 10, 585–590.

Tan, M., and Le, Q. V. (2019). Efficientnet: Rethinking model scaling for convolutional neural networks. arXiv 1905.11946.

Wäldchen, J., and Mäder, P. (2018). Machine learning for image based species identification. Methods Ecol. Evol. 9, 2216–2225.

Wang, Y., Yao, Q., Kwok, J., and Ni, L. M. (2020). Generalizing from a few examples: a survey on few-shot learning. ACM Comput. Surv. 53, 1–34.

Warren, D. L., Glor, R. E., and Turelli, M. (2010). ENMTools: a toolbox for comparative studies of environmental niche models. Ecography 33, 607–611.

Weinstein, B. G. (2018). A computer vision for animal ecology. J. Animal Ecol. 87, 533–545.

Willi, M., Pitman, R. T., Cardoso, A. W., Locke, C., Swanson, A., Boyer, A., Veldthuis, M., and Fortson, L. (2019). Identifying animal species in camera trap images using deep learning and citizen science. Methods Ecol. Evol. 10, 80–91.

Whytock, R.C., Świezewski, J., Zwerts, J.A., Bara-Słupski, T., Pambo, A.F.K., Rogala, M., Bahaa-el-din, L., Boekee, K., Brittain, S., Cardoso, A.W., Henschel, P., Lehmann, D., Momboua, B., Opepa, C.K., Orbell, C., Pitman, R.T., Robinson, H.S., and Abernethy, K.A. (2021). Robust ecological analysis of camera trap data labelled by a machine learning model. Methods Ecol. Evol. 12, 1080–1092.

Yosinski, J., Clune, J., Bengio, Y., and Lipson, H. (2014). How transferable are features in deep neural networks? In Advances in neural information processing systems, Z. Ghahramani, M. Welling, C. Cortes, N. D. Lawrence, and K. Q. Weinberger, Eds. (Curran Associates, Inc), pp. 2220–3328.

Zeiler, M. D., and Fergus. (2014). Visualizing and understanding convolutional networks. Notes Comut. Sci. 8689, 818–833.

